# Defining transcriptomic niches in human fibrotic lung with multi-sample spatial transcriptomics analysis using the MAPLE algorithm

**DOI:** 10.1101/2022.02.28.482296

**Authors:** Hyeongseon Jeon, Carter Allen, Natalia Vanegas-Avendano, Juan Xie, Ahmed Hamdy Ghobashi, Jhonny Rodriguez-Lopez, Lorena Rosas, Cankun Wang, Hao Cheng, Yuzhou Chang, Konstantin Shilo, Anjun Ma, Ana L. Mora, Mauricio Rojas, Qin Ma, Dongjun Chung

## Abstract

Idiopathic pulmonary fibrosis (IPF) is an age-related lung disease marked by patchy parenchymal fibrosis that disrupts gas exchange and leads to respiratory failure. Single-cell RNA-seq studies have revealed cellular heterogeneity in IPF, but the spatial contexts underlying alveolar epithelial cell loss, honeycomb cyst formation, and fibroblastic foci remain unclear, motivating comparisons based on multi-sample spatial transcriptomics (ST) across clinical and anatomical conditions. Here, we present a comprehensive pipeline for analyzing multi-sample lung ST data that accounts for experimental design, featuring the MAPLE algorithm to jointly identify cell-spot subpopulations and discover spatial biomarkers, with validation using 10X Xenium ST data. Our analysis reveals that IPF progression involves epithelial-to-mesenchymal state transitions and region-specific fibrotic programs, characterized by extracellular matrix remodeling in the upper lobes and B-cell-enriched immune niches in the lower lobes. We identify fibrotic regions with senescent cell (SnC) enrichment and demonstrate their spatial colocalization with aberrant basaloid niches.

**Author Summary:** Idiopathic pulmonary fibrosis is a progressive lung disease in which normal lung tissue is replaced by scar tissue in uneven and patchy patterns. Understanding these patterns requires methods that compare multiple tissue samples while preserving information about where genes are expressed. We developed and applied MAPLE, a statistical framework for jointly analyzing multiple spatial transcriptomics samples while accounting for differences between clinical groups and spatial relationships within each tissue section. Using lung tissues from healthy donors and patients with idiopathic pulmonary fibrosis, we identified shared tissue regions across samples and linked them to cell composition, gene expression, and pathological features. We observed a stepwise decrease in alveolar epithelial and endothelial cells across the analyzed conditions, an increase in mesenchymal states, extracellular matrix remodeling in upper-lobe fibrotic regions, and B-cell-associated programs in lower-lobe regions. We also found that senescence-associated signals were spatially concentrated near aberrant basaloid cell niches. These results demonstrate how multi-sample spatial analysis can reveal disease-associated tissue organization that may be missed when tissue samples are analyzed separately. MAPLE provides an interpretable framework for studying spatially heterogeneous diseases across multiple patients and anatomical regions.

## 1. Introduction

Idiopathic pulmonary fibrosis (IPF) is a chronic, progressive, age-related lung disease of unknown origin, characterized by the accumulation of senescent cells (SnCs). IPF leads to excessive extracellular matrix production and lung scarring (fibrosis), which hampers breathing and ultimately causes respiratory failure and death [1]. IPF usually affects individuals over 60 years old, and despite ongoing therapeutic efforts, treatment options remain limited. The median survival time after diagnosis for individuals with IPF is 3-5 years [2]. Given IPF’s poor prognosis and limited treatment options, gaining a deeper understanding of the disease pathogenesis is a top priority for ongoing research.

IPF is highly heterogeneous, with fibrotic lesions interspersed with areas of normal lung parenchyma [3], including spatial heterogeneity across lung lobes that corresponds to distinct diagnostic, functional, and prognostic contexts [4, 5]. To better understand the complexity, single-cell RNA sequencing (scRNA-seq) studies have characterized diverse lung cell subpopulations in IPF lungs, including specialized epithelial, endothelial, mesenchymal, and immune cells. These studies have provided significant insights into disease pathogenesis by identifying altered gene expression and abnormal activation of various signaling pathways [6-14]. Parallel scRNA-seq analyses of normal lung tissues have further enhanced our understanding of the cellular composition under non-disease clinical conditions [15-17]. However, a significant limitation of scRNA-seq studies is the loss of spatial context due to tissue dissociation, which prevents the study of localized cell-cell interactions that likely contribute to the patchy, regionally heterogeneous fibrotic lesions observed in IPF.

Unlike scRNA-seq, spatial transcriptomics (ST) enables gene expression profiling within intact tissue sections, thereby preserving the native spatial organization of cells and their microenvironment in normal and diseased lung tissue. This advancement is critical because spatial proximity and local cellular microenvironments play a key role in determining cellular phenotype, cell state, and tissue function [18]. Commercial ST platforms, such as 10X Genomics Visium, enable transcriptome-wide profiling at spatially defined spots, known as capture spots, each of which collects transcripts from multiple neighboring cells. Recent studies of IPF lungs using ST have revealed spatially restricted epithelial states and region-specific immune cell distributions, shedding light on the local microenvironments that promote fibrosis [19-22] and demonstrating the power of this technology to reveal spatial heterogeneity as a critical aspect of IPF’s complex pathology.

A fundamental objective in ST data analysis is to identify spatial architecture, which refers to the spatial organization of cellular and molecular features within tissue [23]. An essential step toward this goal is detecting cell-spot subpopulations, defined as groups of spatially adjacent spots with similar gene expression profiles. This is usually achieved through clustering that integrates gene expression and spatial coordinates, followed by biological annotation of cellular composition or histological context. Existing methods, such as BayesSpace [24], SPRUCE [25], STAGATE [26], SpaGCN [27], stLearn [28], RESEPT [29], Seurat [30], and Giotto [31], perform well for single tissue sections. However, IPF studies often involve multiple samples from different patients, anatomical regions (e.g., upper vs. lower lobes), or clinical states (e.g., healthy vs. diseased). This creates a need for multi-sample ST clustering methods that jointly capture spatial architecture while modeling dependencies both within and between samples. Such methods are essential to reveal shared and sample-specific spatial patterns that help explain IPF mechanisms. Recently, multi-sample analysis of ST data has gained attention, and several methods have been proposed, including PASTE [32], PASTE2 [33], BASS [34], SPIRAL [35], and PRECAST [36]. However, existing methods remain limited in their ability to accommodate complex experimental designs in multi-sample ST data, as they often lack frameworks for directly modeling how experimental covariates influence tissue architecture probabilities within spatial models. This limitation is particularly problematic for complex tissues such as the lung, where spatial dependencies vary systematically across anatomical regions and clinical conditions.

To address limitations of existing ST clustering methods, we developed MAPLE (Multi-sAmple sPatiaL transcriptomics modEl) as one component of a comprehensive pipeline for analyzing ST data. MAPLE provides a statistically principled framework for jointly identifying cell spot sub-populations across multiple ST data by combining Bayesian mixture modeling of spatial dependencies with logit-linked modeling of covariate effects on spatial domain probabilities, enabling robust and interpretable inference. Building on this, our pipeline integrates multiple steps beyond clustering, including a structured ST experimental design with samples from healthy tissue, lower lobe IPF, and upper lobe IPF; batch correction to align data across samples; annotation of pulmonary regions using markers identified in scRNA-seq studies; identification of region-specific marker genes; and cell-type deconvolution to estimate the cellular composition of each sub-population. By combining these components, we deliver a flexible and interpretable framework for multi-sample lung ST analysis that advances both methodological rigor and biological insight into IPF.

## 2. Results

### Overview: Multi-sample Spatial Transcriptomics Data Analysis Pipeline

Our comprehensive pipeline for analyzing multi-sample lung ST data (**Fig. 1**) integrates both experimental design and analytical procedures, including the MAPLE clustering algorithm and downstream analyses, followed by experimental validation. The experimental design includes age-matched healthy controls (lower lobe sections) and IPF donors (lower and upper lobe sections), with two tissue samples per group (hereafter referred to as ‘samples’). The samples were formalin-fixed and paraffin-embedded (FFPE), and ST data were generated using the 10x Genomics Visium platform, which offers broader transcriptome coverage compared to current cell-level-resolution ST technologies such as the 10X Xenium platform.

**Fig. 1.**
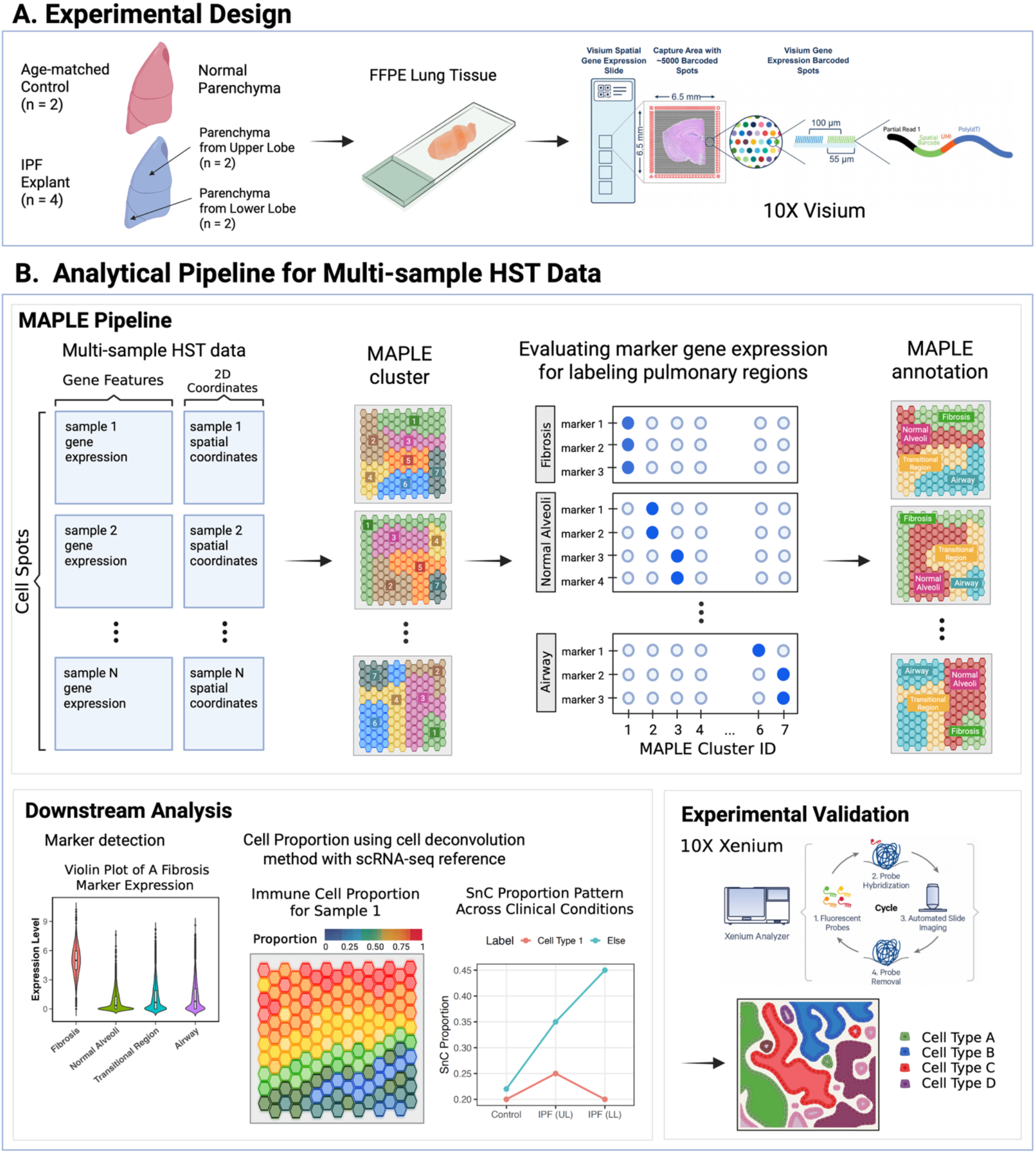
Overview of experimental design and analytical framework for multi-sample lung ST data. (**A**) Experimental protocol and sample processing for multi-sample lung ST data. (**B**) Analytical pipeline including MAPLE clustering, cluster annotation, and downstream analyses, including marker detection, cell-type deconvolution, inference of cell types associated with SnC enrichment, and experimental validation.

To analyze the 10X Visium ST data, we performed batch correction using Seurat’s integration pipeline [37]. Then, we applied our MAPLE algorithm to jointly cluster cell spots across six samples, representing two biological replicates per clinical condition. Annotation of MAPLE-derived clusters was guided by canonical scRNA-seq markers and focused on broader pulmonary regions to ensure robust and interpretable labeling, a strategy particularly suitable for IPF data, which exhibits strong region-specific pathology. The MAPLE-derived cell spot sub-populations were further validated using pathological annotations from H&E-stained images. Then, region-specific biomarkers were identified, improving our understanding of spatial heterogeneity in both healthy and fibrotic lung tissues. Finally, the region-specific biomarkers discovered with the 10X Visium ST were validated using the 10X Xenium ST, which provides single-cell-level resolution, by confirming that their spatial patterns are preserved at the cellular level.

### MAPLE Clustering and Pulmonary Region-Based Annotation

To characterize the different pulmonary regions in multi-sample lung ST data, we applied the MAPLE algorithm to cluster and annotate cell spots using regional markers together. Following a default Seurat pipeline, data integration was performed to remove batch effects across six samples. Dimensionality reduction was conducted using Principal

Component Analysis (PCA), and the first 10 principal components (PCs) were selected based on the elbow method from the explained variance plot. The MAPLE clustering algorithm was then applied to the PCA-transformed feature matrix, with sample-specific clinical conditions included as covariates to capture condition-specific variation in cluster proportions. This choice was motivated by the potential differences in heterogeneity in IPF lung tissue, given its more complex tissue architecture compared with control samples. The optimal number of clusters was determined using the Bayesian Information Criterion (BIC), resulting in eleven clusters that captured distinct underlying patterns in the lung tissue. MAPLE clusters were validated by confirming clear cluster-specific gene expression patterns in the t-SNE embedding (**Fig. S1**).

Cluster annotation was guided by regional canonical markers (**Fig. 2A**), allowing for the assignment of each MAPLE cluster to its corresponding pulmonary region and thereby enabling region-specific analysis of spatial heterogeneity. The spatial distribution of these annotated regions (**Fig. 2B)** reveals how fibrotic, normal parenchymal, immune-rich, airway-associated, and vascular regions are organized across the lung tissue sections. Histological sections originated from same tissues show minimal interstitial involvement in age-matched control lungs, whereas idiopathic pulmonary fibrosis (IPF) samples exhibit extensive remodeling. IPF upper and lower lobes display dense hyalinized fibrous scarring and fibroblast foci, with scattered inflammatory aggregates, highlighting the heterogeneity of fibrotic and inflammatory changes across lobes (**Fig. S2**).

**Fig. 2.**
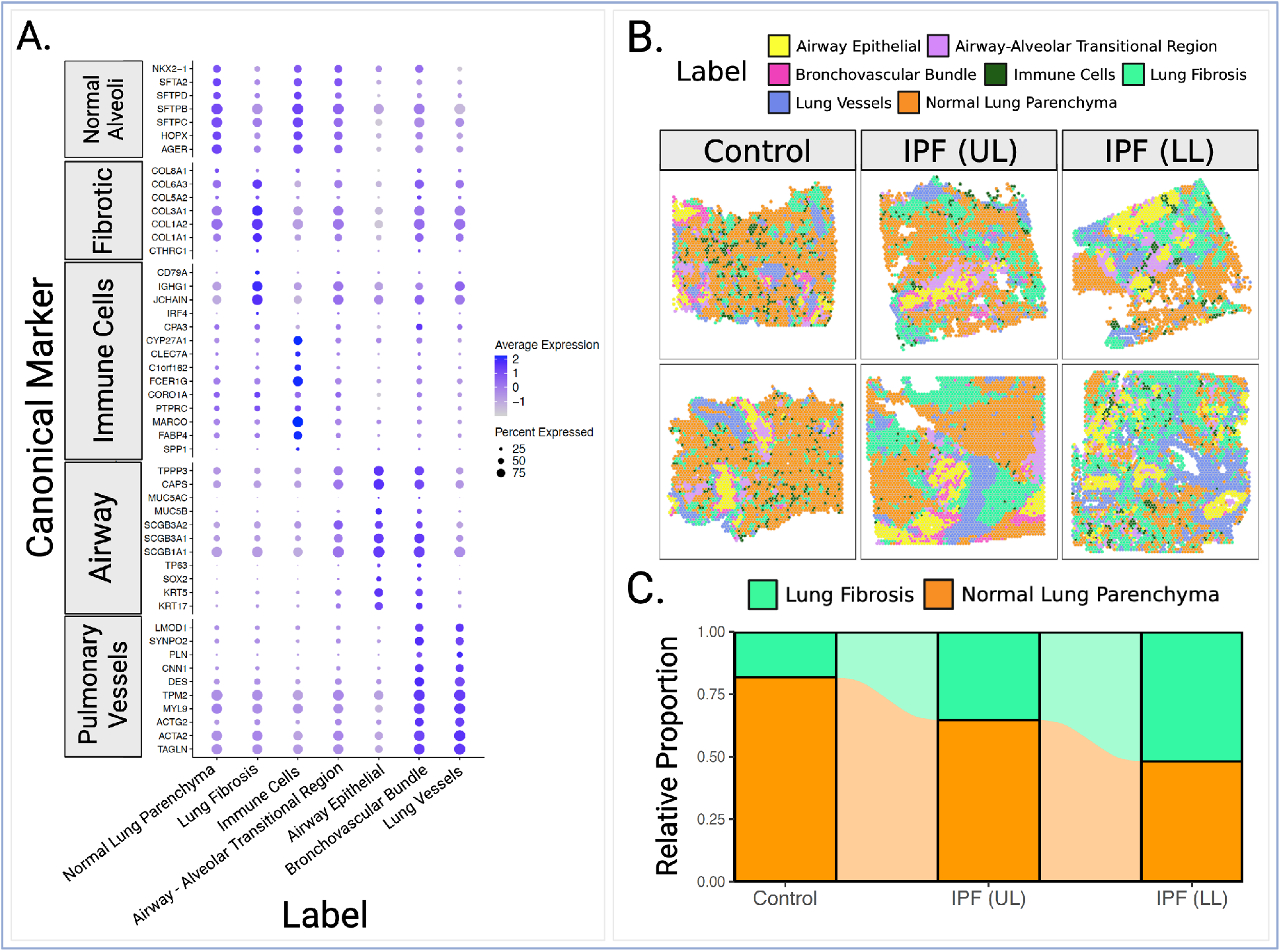
The MAPLE pipeline is visualized with three subfigures. (**A**) Dot plot showing canonical marker gene expression across MAPLE-defined pulmonary regions. (**B**) Visualization of the seven MAPLE-derived regions. (**C**) Alluvial plot illustrating relative changes between lung fibrosis and normal parenchyma across the three clinical conditions.

For downstream analyses, lung fibrosis and normal lung parenchyma served as the primary regions of interest. Fibrotic regions were identified by strong expression of collagen and other ECM-related genes (e.g., *COL1A1, COL1A2, COL3A1, CTHRC1*) and annotated by lung fibrosis. In contrast, clusters characterized by low collagen and immune marker expression and high expression of normal alveolar markers were annotated as normal lung parenchyma. Notably, several immune-related genes, including *CD79A, IGHG1*, and *JCHAIN*, also exhibited elevated expression within the lung fibrosis. Immune cell-rich clusters, defined by elevated lymphoid and myeloid signatures and high expression of normal alveolar markers, were annotated as immune cells. As expected in IPF lung tissue, these immune cell-rich regions are frequently localized near fibrotic regions, consistent with known patterns of immune cell accumulation adjacent to fibrotic lesions in IPF lungs [38]. The final MAPLE annotation (**Fig. 2C**) highlights a progressive increase in fibrotic regions relative to normal parenchyma, moving from controls to IPF upper-lobe tissues and then to IPF lower-lobe tissues.

### Progressive Shift from Alveolar Epithelial to Mesenchymal States Defines IPF Progression

Because each cell spot in the 10x Visium ST data captures transcripts from more than one cell, estimating the underlying cell-type composition is essential for downstream analyses. Accurate estimates of cell-type proportions enable higher-resolution biological interpretation, including the assessment of spatial localization between cell states and the identification of condition-specific trends across pulmonary regions. To infer these cell proportions, we employed TACCO [39], a robust label-transfer framework that maps scRNA-seq-derived cell-type annotations onto spatial transcriptomic spots. By using matched scRNA-seq reference and 10X Visium data derived from the same donors in our analysis, we ensured reliable cell-type proportion estimates with TACCO while preserving the biological relationships present in the original samples. This approach allows us to characterize spatial cell-type distributions in ways that would not be accessible from spatial data alone, thereby enhancing the interpretability and biological relevance of our downstream analyses.

Using these TACCO-derived cell proportion estimates, we observed distinct shifts in cell-type composition across clinical conditions. **Fig. 3A** shows a marked increase in immune cell proportions in IPF, a progressive decline in endothelial and alveolar epithelial cells from healthy tissue to UL and LL IPF samples, and a distinct mesenchymal cell pattern characterized by a peak in UL followed by a slight decrease in LL. When directly comparing mesenchymal and alveolar epithelial proportions, the fraction of mesenchymal cells exceeding alveolar epithelial cells increased from control to UL, as illustrated in **Fig. 3B**. They rose further in LL, reflecting an expanding dominance of mesenchymal states with disease progression. These spatial patterns are also evident in **Fig. 3C-F**, where the color gradients visually capture the expansion of mesenchymal regions and the loss of endothelial signal across IPF samples.

**Fig. 3.**
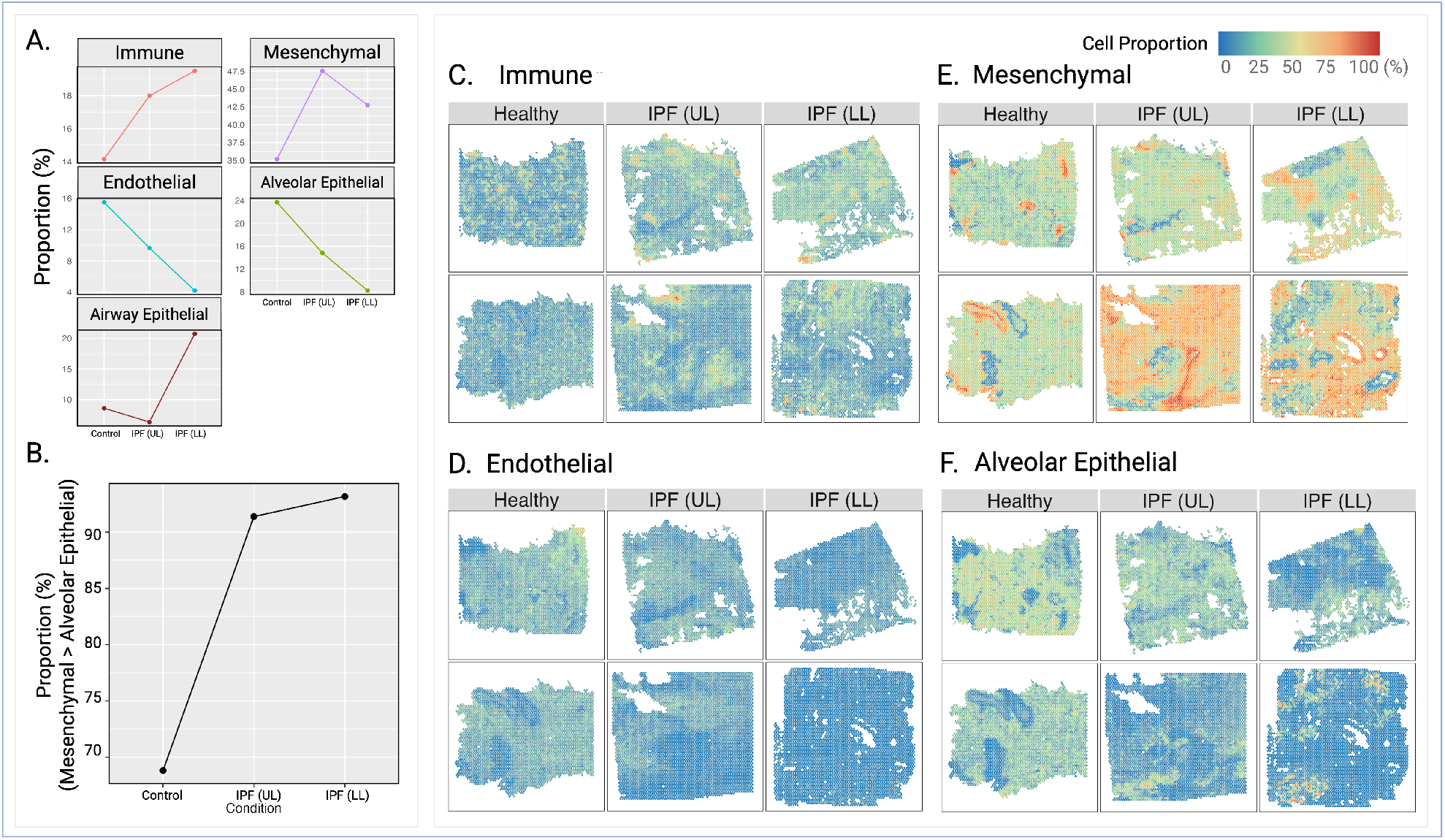
Cell-type deconvolution results and spatial patterns of major cell types. (**A**) Mean cell-type proportions across clinical conditions for immune, mesenchymal, endothelial, alveolar epithelial, and airway epithelial cells. (**B**) Comparison of mesenchymal and alveolar epithelial proportions, showing the extent to which mesenchymal proportions exceed epithelial proportions. (**C-F**) Spatial feature plots showing the distribution of immune (**C**), endothelial (**D**), mesenchymal (**E**), and alveolar epithelial (**F**) cell proportions across Healthy, UL IPF, and LL IPF tissue sections.

### Region-Specific Fibrotic Programs in IPF: ECM Remodeling in Upper Lobes and B-Cell Enrichment in Lower Lobes

As part of our downstream analysis, we conducted spatially informed transcriptomic comparisons by integrating MAPLE-defined regions with pathological annotations to identify markers associated with IPF-specific pathological states. We focused on contrasting healthy normal parenchyma (Normal PA) with two anatomically distinct fibrotic regions observed in IPF: upper-lobe fibrotic foci (IPF UL_FF) and lower-lobe fibrotic foci (IPF LL_FF). Using Seurat’s ‘FindMarkers’ function (fold change of 8, adjusted p-value BH < 0.05), we identified 199 genes upregulated in IPF UL_FF and 92 genes upregulated in IPF LL_FF relative to Normal PA. Pathway enrichment analysis using ToppFun [40] revealed that UL_FF was enriched for extracellular matrix (ECM) remodeling and interferon signaling pathways, whereas LL_FF showed strong enrichment for B-cell-associated pathways, including B-cell receptor (BCR) signaling and CD22-mediated BCR regulation, along with robust upregulation of canonical B-cell markers such as *CD79A, MZB1*, and *POU2AF1*.

To experimentally validate these region-specific markers at single-cell resolution, we employed 10X Xenium, which enables targeted gene profiling with preserved spatial context. Xenium imaging demonstrated that *CD79A*^+^, *MZB1*^+^, and *POU2AF1*^+^ B-cells were spatially localized around *CTHRC1*^+^ fibroblastic foci, particularly within LL_FF samples, confirming the spatial co-occurrence of B-cell programs within fibrotic niches. This spatial pattern supports a potential immune-stromal interaction and highlights distinct molecular programs driving upper-versus lower-lobe fibrotic lesions in IPF (**Fig. 4**).

**Fig. 4.**
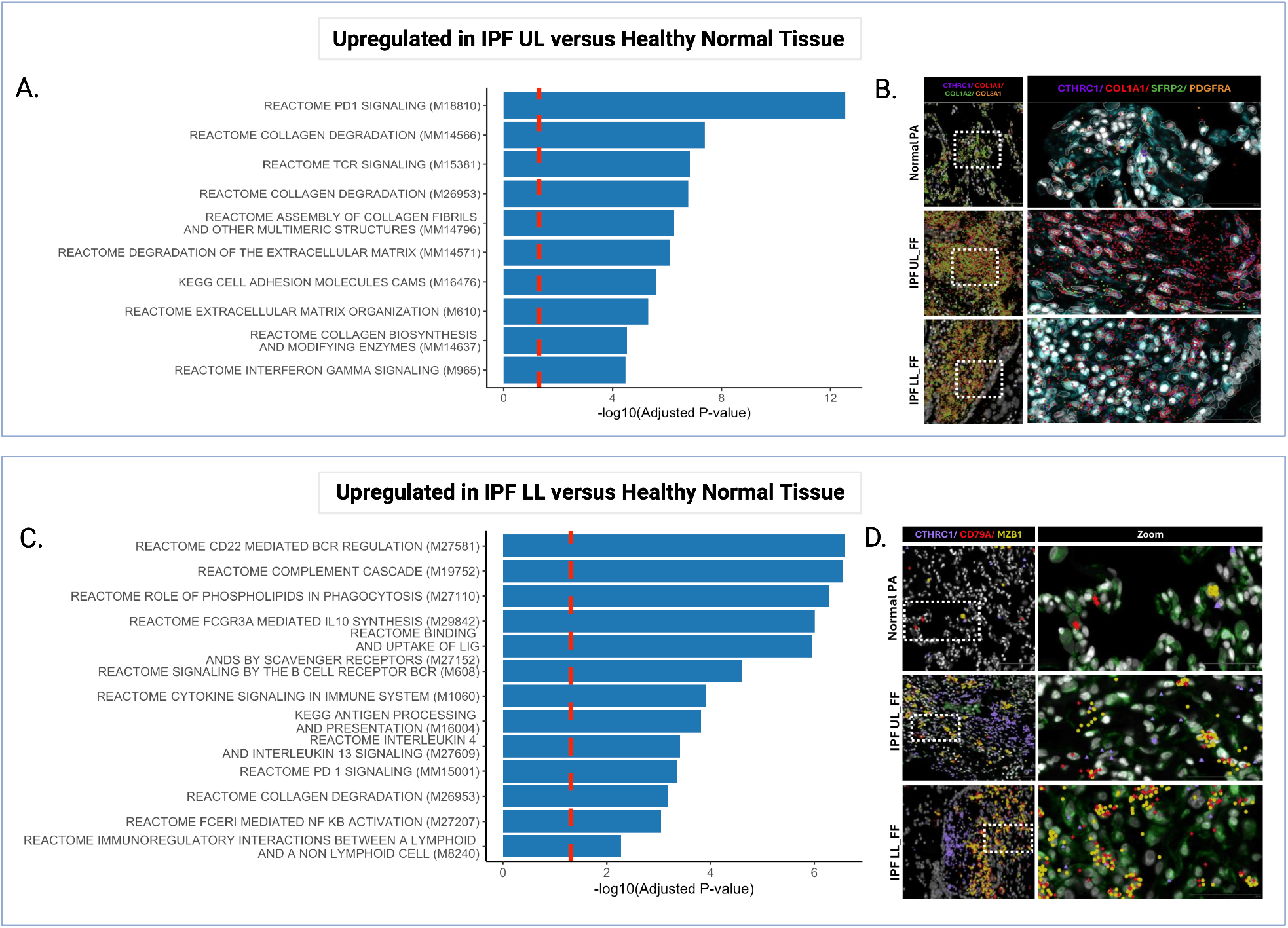
(**A**) Top pathways enriched in genes upregulated in UL IPF versus healthy parenchyma. (**B**) 10X Xenium-based validation of UL-enriched markers, showing increased expression in fibrotic regions. (**C**) Top pathways enriched in genes upregulated in LL IPF versus healthy parenchyma. (**D**) 10X Xenium-based validation of LL-enriched markers, confirming elevated expression in IPF tissue.

To experimentally validate these region-specific markers at single-cell resolution, we employed 10X Xenium, a ST platform that enables targeted gene profiling. We focused on fibrotic foci and their surrounding regions as regions of interest and quantified the proportion of *CD79A*^+^, *MZB1*^+^, and *POU2AF1*^+^ cells as markers of B cells, using age-matched parenchyma as a control.

Our data showed, as expected, that *CTHRC1*^+^ cells were highly enriched within fibrotic foci. In contrast, B cells were significantly enriched in the regions surrounding these foci and were largely absent from age-matched parenchyma. These findings indicate that B cells are spatially localized around *CTHRC1*^+^ fibroblastic foci, confirming the spatial co-occurrence of B cell transcriptional programs within fibrotic niches. This spatial organization supports the presence of potential immune-stromal interactions and highlights distinct molecular programs operating within fibrotic lesions in IPF. These analyses and results are summarized in **Figs. 4B and D** and **Fig. S3**.

### Spatial Colocalization of Senescent Cells and Aberrant Basaloid Niches in IPF Lung Fibrosis

We performed additional downstream analyses to identify SnC-enriched cell spots by integrating MAPLE annotations and curated senescence markers from independent databases. To construct a high-confidence SnC gene panel, we first aggregated senescence-associated genes from multiple sources [41-46], including SenMayo [42], FRIDMAN [43], and CellAge [41], which are listed in **Table S1**. From the initial list of senescence-related genes, we selected genes that appeared in at least three databases, yielding a robust consensus set of 161 markers. The list was further refined by selecting genes exhibiting elevated expression in fibrotic regions, resulting in a final panel of 11 IPF-relevant SnC markers: *TNF, SERPINE1, WNT2, MMP13, IGF1, MMP14, FGF7, MMP2, IGFBP4, MMP9*, and *GDF15*. (**Fig. 5D**).

**Fig. 5.**
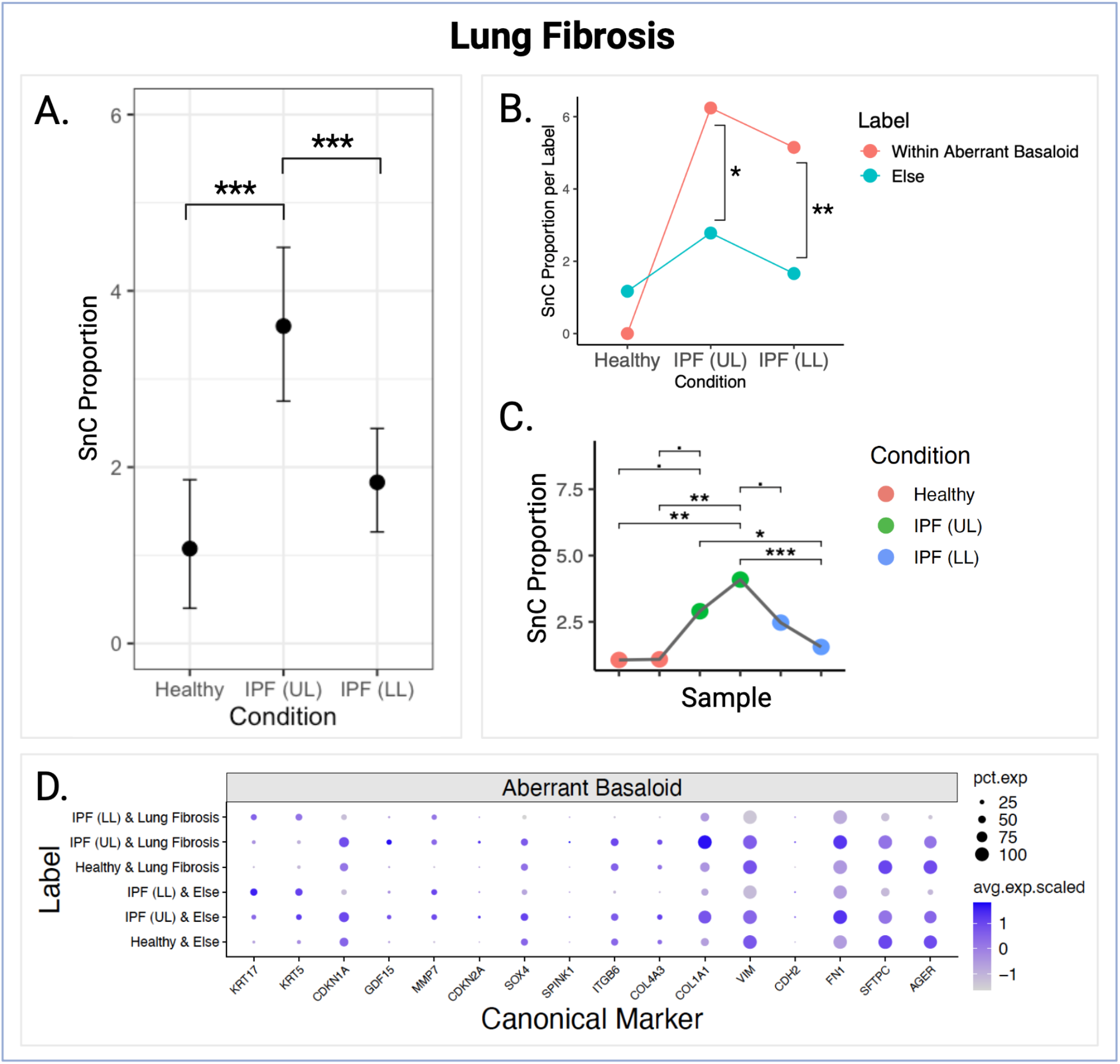
SnC enrichment patterns and their spatial association with aberrant basaloid regions. (**A**) Proportion of SnC-enriched spots across Healthy, IPF (UL), and IPF (LL) lung fibrosis regions. (**B**) SnC proportion within and outside aberrant basaloid regions across conditions, showing significant colocalization between SnC-enriched and aberrant basaloid-enriched spots in IPF conditions. (**C**) Sample-level SnC proportions across conditions, demonstrating a consistent increase in IPF samples. (**D**) Dot plot showing canonical aberrant basaloid marker expression across spatial regions, highlighting strong marker expression within SnC-enriched areas.

The stepwise selection of the genes, including the initial conservatively curated SnC marker list, the subsequent intersection with Seurat-derived feature genes, and the final fibrosis-enriched subset, is summarized in **Table S1**.

Using the average expression of these selected 11 genes, we quantified SnC enrichment across spatial cell spots and identified SnC-enriched spots in a conservative manner, by applying a Gaussian mixture model and defining enriched spots as those exceeding the 95th percentile of the higher-expression component. In parallel, cell-type-enriched cell spots were determined using cell-type proportions with a histogram-based thresholding approach. After deriving these enrichment maps, we estimated SnC proportions within lung fibrosis regions and observed that SnCs were consistently more abundant in IPF than in healthy tissue, a pattern that remained stable across individual samples (**Figs. 5A and C**). To investigate the basis of this increase, we assessed the spatial relationship between SnC-enriched cell spots and aberrant basaloid-enriched cell spots. As shown in **Fig. 5B**, IPF samples exhibited significant colocalization between SnCs and aberrant basaloid, indicating that the elevated SnC burden in fibrotic lesions is primarily driven by their spatial proximity to aberrant basaloid cells. This interpretation was further supported by marker expression patterns: canonical basaloid markers (*KRT17, COL17A1, MMP7*) were strongly expressed within SnC-enriched cell spots (**Fig. 5D**). Together, these findings suggest that SnC accumulation in IPF fibrosis is closely linked to aberrant basaloid expansion, and that localized senescence within basaloid niches may contribute to the distinct SnC patterns observed across UL and LL regions.

## 3. Discussion

Our study presents a statistically robust multi-sample ST analysis pipeline centered around the MAPLE clustering method, which effectively models spatial dependence by capturing the varying probabilities of anatomical cluster membership across distinct clinical conditions. This pipeline integrates key components, from experimental design to annotation validation, to support interpretable and biologically grounded analysis. Specifically, we used MAPLE to generate spatial annotations that reflect anatomically and pathologically meaningful pulmonary regions. These annotations were independently validated using cell type deconvolution via TACCO and single-cell resolution 10X Xenium ST data. Beyond validation, we demonstrated how MAPLE-derived pulmonary regions can guide downstream analyses in IPF. We performed differential expression analyses to identify region-specific gene markers and validated these findings in Xenium ST data. This integrative approach enabled us to spatially contextualize immune activation and senescence-associated signatures, thereby providing a practical and extensible framework for applying MAPLE annotations to investigate disease mechanisms in complex tissues, such as the fibrotic lung.

Our integrative analysis pipeline combining 10X Visium ST, Xenium ST, and scRNA-seq reveals a spatial and functional convergence between immune activation, particularly B-cell-driven responses, and cellular senescence in immune cell-enriched regions. Pathway analysis of differentially expressed genes upregulated in IPF LL compared to healthy controls highlighted B-cell-related signaling, which was further validated at single-cell resolution using Xenium ST. In IPF UL, fibrotic lesions co-localized with B cells, whereas in IPF LL, B-cell-rich areas appeared spatially distinct from fibrotic lesions marked by *CTHRC1*, suggesting region-specific immune architecture.

ST analysis using 10X Visium revealed that SnC-enriched spots were preferentially located within immune cell-rich regions, particularly in IPF LL. Interestingly, SnC accumulation was relatively lower in the fibrotic areas in the lower lobe, suggesting that senescent cells are spatially enriched outside the fibrotic core. This pattern is consistent with our single-cell spatial validation, which showed that B cells are spatially separated from fibrotic lesions in IPF LL. While we did not directly test co-expression between SnC and immune markers, the shared localization of SnCs and immune cells, distinct from fibrotic areas, raises the possibility of an interaction between immune senescence and fibrotic niches. These findings support a model where chronic immune activation and senescence-associated remodeling may converge in specific regions, potentially acting in parallel with, but spatially distinct from, fibrosis. Future studies should explore whether immune-mediated senescence contributes to broader tissue dysfunction or represents a byproduct of chronic inflammation in adjacent immune microenvironments.

While the current framework provides a strong foundation for multi-sample ST analysis, future studies could further refine spatial domain identification by introducing joint clustering-annotation models that integrate ST data with histological and clinical features in a unified probabilistic manner. Such an approach would move beyond the current two-step process of clustering followed by manual annotation, and allow for more biologically informed spatial domain identification. Additionally, nonparametric or hierarchical clustering models could be explored to relax the assumption of a fixed number of clusters, better reflecting the cellular heterogeneity within each ST spot. For senescence analysis, future studies should functionally subclassify SnC markers, such as those involved in SASP, DNA damage, or cell cycle arrest, to capture the diversity of senescent programs and their spatial associations. Finally, incorporating time-resolved or pseudotime modeling into ST analyses may help uncover the dynamic interplay between immune activation, senescence, and fibrotic progression across disease stages.

## 4. Materials and Methods

### 4.1 Ethics Statement

Collection of human lung tissue was approved by the Institutional Review Board (IRB) of The Ohio State University (OSU) under study ID 2021H0180. Tissue collection was performed by the Comprehensive Transplant Center (CTC) Human Tissue Biorepository at OSU with informed consent and research authorization obtained from all donors under the Total Transplant Care Protocol. The CTC operates in accordance with NCI and ISBER Best Practices for Repositories. This study constitutes secondary research, and the study team did not participate in participant recruitment or any participant-facing activities.

### 4.2 Sample preparation and demographics information

#### 4.2.1 Preparation of FFPE-lung tissue samples

The demographic summary of the study population is shown in **Table 1**. Lung tissues were collected from the upper and lower lobes of lung explants of patients diagnosed with idiopathic pulmonary fibrosis (IPF) and from the parenchymal anatomical region of age-matched normal donors. The lung tissue was minced into small pieces and fixed for 24 h in 10% neutral buffered formalin, and then transferred to 70% ethanol and embedded in paraffin (wax) to create a FFPE lung tissue block.

**Table 1.**
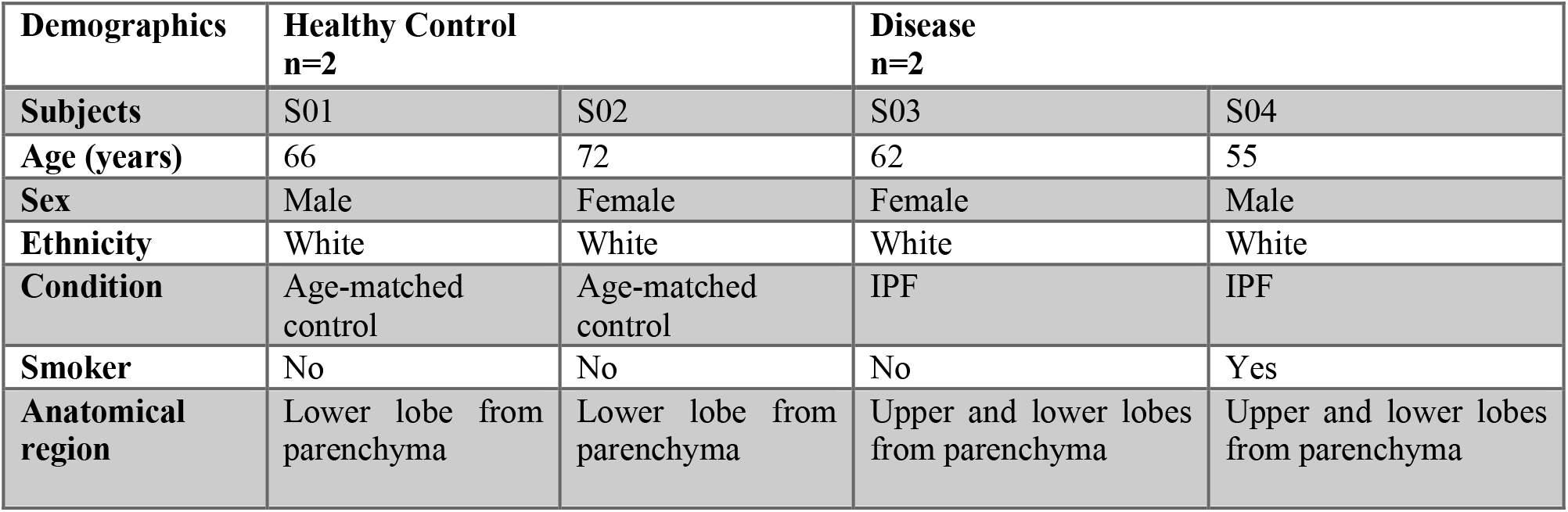
Demographic characteristics of the study subjects.

#### 4.2.2 Visium Spatial Gene Expression from FFPE

Tissue sections of ≤ 6.5 × 6.5 mm at 5 µm were then cut from the FFPE blocks and placed onto Visium Spatial Slides (10X Genomics) according to the tissue preparation guide. The tissue sections onto Visium Spatial Slides were deparaffinized and stained with Hematoxylin & Eosin, and then scanning was performed with EVOSTM M7000 microscope (Invitrogen) using a 10X objective. FFPE libraries were prepared according to the 10X Genomics Guide, and sequenced on an Illumina sequencer to a read depth of a least 25k reads/spot, with read lengths of 28 cycles Read 1, 10 cycles i7 index, 10 cycles i5 index, and 50 cycles Read 2S. 10X Visium ST data were obtained using Space Ranger 1.1.0 from 10X Genomics (https://support.10xgenomics.com). The libraries were mapped to an Ensembl 93-based reference (10X-provided GRCh38 reference, version 2.0.0). For each slide, gene expression in the Spatial assay was normalized using SCTransform, and quality-control metrics were computed, including the number of detected genes and UMIs per spot, as well as the percentage of mitochondrial and hemoglobin genes. SCTransform-normalized objects were then combined and split by sample, and Seurat’s SCT-based integration workflow was applied, with 3,000 integration features selected across samples. Data were prepared using PrepSCTIntegration, integration anchors were identified with FindIntegrationAnchors, and an integrated expression matrix was obtained with IntegrateData, utilizing SCT normalization. The integration analysis was performed with Seurat v4.1.1 in R v4.2.1.

#### 4.2.3. Single-cell RNA-Seq from FFPE

Fixed RNA Profiling was performed on parenchymal lung tissue from two age-matched normal donors and on upper and lower lobe samples from two distinct IPF donors, corresponding to the same donors and sections analyzed by Visium spatial transcriptomics. For each sample, two tissue sections, each 50 µm thick, were then cut from the FFPE blocks and placed into a 1.5ml tube. Then, cells isolated from FFPE tissue were used for Fixed RNA Profiling and library preparation (10X Genomics), followed by next-generation sequencing at the Advanced Genomics Core at the University of Michigan. Gene expression libraries were sequenced on an Illumina sequencer with read lengths of 28 cycles Read 1, 10 cycles i7 index, 10 cycles i5 index, 90 cycles Read 2. Single-cell RNA sequencing data were first demultiplexed, aligned, and quantified using the Cell Ranger pipeline (version 7.1.0; 10x Genomics) according to the manufacturer’s recommendations. Reads were mapped to the GRCh38 reference (refdata-gex-GRCh38-2020-A), and transcript annotation was augmented with the Chromium Human Transcriptome Probe Set v1.0.1 (GRCh38). Count matrices were imported into R (version 4.2.3) using Seurat (version 4.4.0). Ambient RNA contamination was identified and removed, and the proportions of mitochondrial, hemoglobin, and ribosomal gene expression were calculated for each cell. Data were then normalized using SCTransform with default parameters, retaining the 3,000 most variable features for dimensionality reduction. Principal component analysis (PCA) was performed, and batch effects were corrected with Seurat’s pipeline. Integrated analysis, along with batch correction, was conducted via Seurat’s IntegrateData function. Finally, neighborhood graphs were constructed on the harmonized principal components (PCs), and clusters were detected using the FindNeighbors and FindClusters functions. *EPCAM* and *CDH1* were used as canonical markers for epithelial cells, *PDGFRA* and *COL1A1* were used as canonical markers for mesenchymal cells, *PECAM1* was used as a canonical marker for endothelial cells, and finally, *PTPRC* was used as a canonical marker for immune cells.

#### 4.2.4 Xenium In Situ

Gene Panel Design. A total of 389 genes were analyzed using Chemistry version v1. Of these, 289 genes were selected from the Xenium human lung gene expression panel by 10X Genomics, while an additional 100 genes were included from a custom-designed panel (CH7CY9) designed by Mora and Rojas Lab. This custom panel was curated based on single-cell transcriptomic data from human lung tissue generated by the Mora and Rojas Lab, with a focus on genes relevant to cell type identification and potential involvement in IPF pathogenesis.

Cell Segmentation. To enhance cell boundary resolution, the Xenium In Situ Cell Segmentation Kit (PN-1000662, 10X Genomics) was used. This kit utilizes a specialized staining protocol, combined with an algorithmic segmentation approach powered by custom-trained machine learning models, which were developed and validated by 10X Genomics.

Sample Preparation and Run. FFPE-lung tissue was sectioned at 5×6 mm at 5µm onto a Xenium slide. The tissue sections analyzed correspond to the same donors and IPF samples previously used for single-cell RNA-seq and Visium ST. Sections were treated to access the RNA for labeling with circularizable DNA probes. Probe ligation generated a circular DNA probe, which was enzymatically amplified. Slides were placed in the Xenium Analyzer (Instrument serial number XETG00239), where the sample was undergoing successive rounds of fluorescent probe hybridization, imaging, and removal, creating bright, easy-to-image signals with a high signal-to-noise ratio. An optical signature specific to each gene was generated, enabling the identification of the target gene. Finally, a spatial map of the transcripts in the entire tissue section was built using Xenium analysis software version 2.0.0.10 and instrument software version 2.0.1.0, which allowed immediate exploration of the assay’s subcellular readout.

Image Generation. The assignment of transcripts to cells, the precision of specific transcript locations, and the verification of cell segmentation were performed in the final step of the analysis workflow using Xenium Explorer v3.0 software by 10X Genomics.

### 4.3 Details of the MAPLE Method

#### 4.3.1 Joint identification of cell sub-populations in multi-sample ST data

We detect spatially informed cell sub-populations in each tissue sample using a Bayesian multivariate finite mixture model with prior distributions specified to induce correlation in mixture component assignments between neighboring cell spots within each sample. First, we let l = 1, …, L index the individual tissue samples in a given spatial transcriptomics data set, where the total number of cell spots sequenced in each sample is denoted n_1_. We index each cell spot in the multi-sample data set as i = 1, …, N, where the total number of cell spots present in the experiment is given by 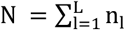. For each cell spot, we denote gene expression features with the length-g vector **y**_**i**_. We assume **y**_**i**_ arises from a K component finite mixture model given by

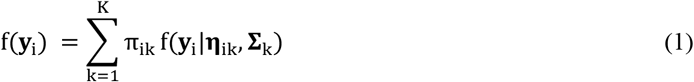

where π_ik_ is a mixing weight that represents the probability of cell spot i belonging to mixture component k, which may itself be modeled as a function of external, f(**y**_i_|**η**_ik_, **∑**_k_) denotes a g-dimensional multivariate normal density with length g location vector **η**_ik_ and g × g variance-covariance matrix **∑**_k_. To allow for spatial heterogeneity in average gene expression profiles within sub-populations, we model **η**_ik_ as

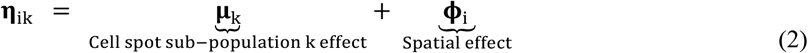

where **µ**_k_ is a length-g mean gene expression profile for mixture component k, and spatial autocorrelation in features among neighboring cell spots is achieved through assuming multivariate conditionally autoregressive (MCAR) priors [47] for the spot-specific random effects **ϕ**_i_. That is, we assume

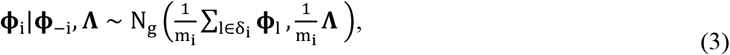

where **ϕ**_−i_ denotes the spatial random effects for all spots except spot i, **Λ** is a g × g variance-covariance matrix for the elements of **ϕ**_i_, m_i_ is the number of neighbors of spot i, and δ_i_ is the set of all neighboring spots to cell spot i. We enforce Cov(**ϕ**_i_, **ϕ**_j_) = 0 when spots i and j are from different tissue samples. As described in [48], we ensure a proper posterior distribution for each **ϕ**_i_ by enforcing a sum-to-zero constraint on the elements of each **ϕ**_i_ for i = 1, …, n. We complete a fully Bayesian model specification by assuming conjugate priors for the mixture component-specific multivariate normal model as **µ**_k_ ∼ N_g_ (**µ**_0k_, **V**_0k_) and **∑**_k_ ∼ IW(***v***_0k_, **S**_.k_). Further, we specify **Λ** ∼ IW(**λ**_0_, **D**_0_). By default, we specify weakly informative priors [49] by setting **µ**_0k_ = **0**_g×1_, **V**_0k_ = **S**_0k_ = **I**_g×g_, and *v*_0k_ = g + 2, which gives E(**∑**_k_) = **I**_g×g_, where **I**_g×g_ is the g × g identity matrix. Similarly, we set λ_0_ = λ_0k_ = g + 2 and **D**_0_ = **D**_0k_ = **I**_g×g_.

Models (1) and (2) feature a number of desirable properties in the context of multi-sample ST data analysis. First, information sharing between samples is achieved by common cell spot sub-population parameters **µ**_k_ and **∑**_k_, thus supporting the utility of integrated multi-sample analysis relative to sample-specific analyses. Relatedly, in addition to inferring cell spot sub-population labels, we may characterize sub-populations using posterior estimates of **µ**_k_ and **∑**_k_, which represent the average gene expression profile and gene-gene correlation of each sub-population, respectively. Additionally, the contribution of each observation **y**_i_ to each cell spot sub-population is governed by the continuous probabilities π_i1_, …, π_iK_. These parameters allows for explanation of cell spot sub-population membership in terms of available covariates. Finally, we note that while we focus this work on modeling continuous features derived from feature mining approaches, our proposed framework may be extended to accommodation of normalized gene expression values directly by allowing for multivariate skew-normal mixture component densities as detailed in [25] and implemented in the R package maple.

To facilitate posterior inference, we introduce the latent cell spot sub-population indicators z_1_, …, z_n_, where z_i_ ∈ {1, …, K} denotes to which cell spot sub-population cell spot i belongs. We assume z_i_ ∼ Categorical(π_i1_, …, π_iK_). We assign cell spots to discrete cell spot sub-populations using the maximum a posteriori estimates 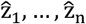. However, unlike existing methods, we accompany these discrete parameter estimates with continuous uncertainty measures to account for (i) the semi-continuous nature of cell type differentiation and (ii) the statistical uncertainty inherent to cell spot sub-population identification.

To quantify the effect of sample-level covariates on tissue architecture in multi-sample ST data, we use an embedded multinomial logit regression model for the spot-level mixture component probabilities *π*_*ik*_:

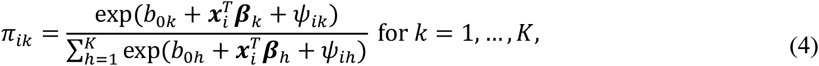

where *b*_0*k*_ is an intercept to adjust for varying cell spot subpopulation sizes, ***x***_*i*_ is a *p*-length vector of covariates specific to spot *i*, ***β***_*k*_ an associated *p*-length vector of regression coefficients for mixture component *k*, and *ψ*_*ik*_ is a spatial random effect allowing spatially-correlated variation with respect to 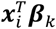. Since specification of a reference category is necessary to ensure an identifiable model formulation in terms of *K* − 1 non-redundant categories, we specify mixture component 1 as the reference category and fix *b*_01_ = 0, ***β***_1_ = **0**_*p*×1_, and *ψ*_*i*1_ = 0 for all *i* = 1, …, *N* accordingly. To introduce spatial association into the component membership model, we assume univariate intrinsic CAR priors for *ψ*_*ik*_:

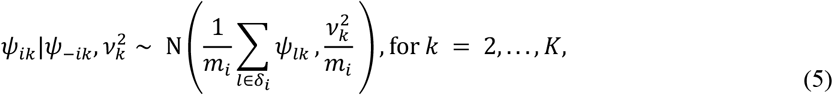

where 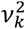 is a mixture component-specific variance for *ψ*_*ik*_, and Cov(*ψ*_*ik*_, *ψ*_*ik*_) = 0 for all *k* = 2, …, *K* when cell spots *i* and *j* are from different samples. This ensures that spatial correlation in mixture component assignments is not introduced between distinct spatial entities (i.e., tissue slices). We assign a conjugate multivariate normal prior for the remaining regression coefficients as 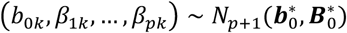, where 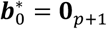, and 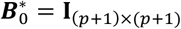, for *k* = 2, …, *K*.

The utility of model (4) lies largely in its generalizability to arbitrary spatial transcriptomics experimental designs, where spot-level covariates ***x***_*i*_ may be specified based on available metadata of a given experiment. By adopting a regression approach, we may assess the effect of experimental covariates such as treatment group or disease status, while adjusting for possible sample-specific confounders like disease stage, sex, age, or batch identifiers.

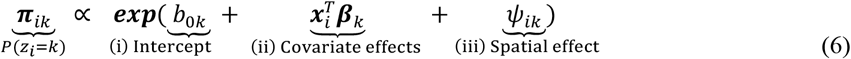

Intuitively, as shown above in equation (6), we construct model (4) to serve three important functions: (i) adjustment for varying cell spot sub-population sizes through the intercept *b*_0*k*_ to account for the fact that we do not necessarily wish for the probability of a randomly selected cell spot belonging to a certain cell spot sub-population to be proportional to the size of that sub-population; (ii) assessment of covariate effects or adjustment for any other confounders of cell spot sub-population membership probability through 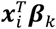; and (iii) the introduction of spatial correlation among neighboring cell spots within the same tissue sample via *ψ*_*ik*_. Critically, by adopting a multinomial regression approach, we avoid the pitfalls of univariate comparison of cell spot sub-population proportions across samples such as unaccounted for negative bias in cell spot sub-population proportions within samples.

#### 4.3.2. Software implementation

MAPLE is freely available as an interactive R package on GitHub at https://github.com/carter-allen/maple, compatible with Mac, Windows, and Linux platforms (GPL >= 2 license). The maple package seamlessly integrates with standard Seurat [30] workflows through a unified modeling interface and interactive visualization functions. The maple package estimates model parameters using efficient Gibbs sampling routines implemented in C++ using Rcpp [50], and interactively visualizes the resultant tissue architecture using the Shiny framework [51]. Run-time for a typical ST data analysis with roughly 5,000 cell spots requires approximately 1 minute per 1,000 iterations on an M1 Apple iMac desktop with 8GB RAM, making it feasible to analyze ST data on a personal computer. After parameter estimation, users may (i) interactively visualize the inferred cell spot labels and annotate cell spot sub-populations, (ii) compute and interactively visualize uncertainty measures that allows assessment of confidence in cell spot labels, and (iii) visualize the relative changes between samples or groups of samples in cell spot sub-population abundances, while accounting for any specified covariates of interest.

## 5. Data Availability

The spatial transcriptomics, single-cell RNA-sequencing, and Xenium datasets supporting this study will be deposited in an appropriate public repository. Repository names and accession numbers are currently pending and will be added when available and no later than acceptance. The data and processed files required for editorial assessment and peer review will be made available to the journal during the review process.

The MAPLE R package is publicly available at https://github.com/carter-allen/maple. The analysis scripts underlying the results and figures presented in this study will be deposited in a public code repository. The repository link and version identifier will be added when available and no later than acceptance.

## 6. Author Contributions

H.J. conceived and led the analytical strategy, performed data and statistical analyses, and led manuscript writing. C.A. developed the core methodology and analytical framework. N.V.-A. led ST sample preparation and experimental validation, and A.H.G. performed validation-related computational analyses. J.X. led the cell-type deconvolution pipeline and its methodological implementation, and C.W. and H.C. contributed to the implementation of cell-type deconvolution analyses. J.R.-L. contributed to initial data processing, generated reference data for cell-type deconvolution, and contributed to pulmonary region annotation. L.R. contributed to experimental validation and data interpretation. K.S. conducted pathological evaluation and validation. A.M. and Y.C. reviewed the manuscript and provided feedback. D.C., Q.M., M.R., and A.L.M. supervised the study and guided the overall research direction. All authors reviewed and approved the final manuscript.

## References

1. Mora, A.L., et al., Emerging therapies for idiopathic pulmonary fibrosis, a progressive age-related disease. Nat Rev Drug Discov, 2017. 16(11): p. 810.

2. Maher, T.M., et al., Global incidence and prevalence of idiopathic pulmonary fibrosis. Respir Res, 2021. 22(1): p. 197.

3. Raghu, G., et al., Idiopathic Pulmonary Fibrosis (an Update) and Progressive Pulmonary Fibrosis in Adults: An Official ATS/ERS/JRS/ALAT Clinical Practice Guideline. Am J Respir Crit Care Med, 2022.205(9): p. e18–e47.

4. Tanaka, Y., et al., Standardised 3D-CT lung volumes for patients with idiopathic pulmonary fibrosis. Respir Res, 2022.23(1): p. 142.

5. Prasad, R., et al., Diagnosis of idiopathic pulmonary fibrosis: Current issues. Intractable Rare Dis Res, 2015. 4(2): p. 65–9.

6. Adams, T.S., et al., Single-cell RNA-seq reveals ectopic and aberrant lung-resident cell populations in idiopathic pulmonary fibrosis. Sci Adv, 2020. 6(28): p. eaba1983.

7. Morse, C., et al., Proliferating SPP1/MERTK-expressing macrophages in idiopathic pulmonary fibrosis. Eur Respir J, 2019. 54(2).

8. Peyser, R., et al., Defining the Activated Fibroblast Population in Lung Fibrosis Using Single-Cell Sequencing. Am J Respir Cell Mol Biol, 2019. 61(1): p. 74–85.

9. Reyfman, P.A., et al., Single-Cell Transcriptomic Analysis of Human Lung Provides Insights into the Pathobiology of Pulmonary Fibrosis. Am J Respir Crit Care Med, 2019. 199(12): p. 1517–1536.

10. Tsukui, T., et al., Collagen-producing lung cell atlas identifies multiple subsets with distinct localization and relevance to fibrosis. Nat Commun, 2020. 11(1): p. 1920.

11. Xu, Y., et al., Single-cell RNA sequencing identifies diverse roles of epithelial cells in idiopathic pulmonary fibrosis. JCI Insight, 2016. 1(20): p. e90558.

12. Jia, M., et al., Early events marking lung fibroblast transition to profibrotic state in idiopathic pulmonary fibrosis. Respir Res, 2023.24(1): p. 116.

13. Habermann, A.C., et al., Single-cell RNA sequencing reveals profibrotic roles of distinct epithelial and mesenchymal lineages in pulmonary fibrosis. Sci Adv, 2020. 6(28): p. eaba1972.

14. Carraro, G., et al., Single-Cell Reconstruction of Human Basal Cell Diversity in Normal and Idiopathic Pulmonary Fibrosis Lungs. Am J Respir Crit Care Med, 2020.202(11): p. 1540–1550.

15. Travaglini, K.J., et al., A molecular cell atlas of the human lung from single-cell RNA sequencing. Nature, 2020. 587(7835): p. 619–625.

16. Sikkema, L., et al., An integrated cell atlas of the lung in health and disease. Nat Med, 2023.29(6): p. 1563–1577.

17. Guo, M., et al., Guided construction of single cell reference for human and mouse lung. Nat Commun, 2023. 14(1): p. 4566.

18. Williams, C.G., et al., An introduction to spatial transcriptomics for biomedical research. Genome Med, 2022. 14(1): p. 68.

19. Vannan, A., et al., Image-based spatial transcriptomics identifies molecular niche dysregulation associated with distal lung remodeling in pulmonary fibrosis. bioRxiv, 2023.

20. Franzén, L., et al., Mapping spatially resolved transcriptomes in human and mouse pulmonary fibrosis. Nat Genet, 2024. 56(8): p. 1725–1736.

21. Eyres, M., et al., Spatially resolved deconvolution of the fibrotic niche in lung fibrosis. Cell Rep, 2022. 40(7): p. 111230.

22. Blumhagen, R.Z., et al., Spatially distinct molecular patterns of gene expression in idiopathic pulmonary fibrosis. Respir Res, 2023.24(1): p. 287.

23. Jeon, H., et al., Statistical Power Analysis for Designing Bulk, Single-Cell, and Spatial Transcriptomics Experiments: Review, Tutorial, and Perspectives. Biomolecules, 2023. 13(2).

24. Zhao, E., et al., Spatial transcriptomics at subspot resolution with BayesSpace. Nat Biotechnol, 2021. 39(11): p. 1375–1384.

25. Allen, C., et al., A Bayesian multivariate mixture model for high throughput spatial transcriptomics. Biometrics, 2022.

26. Dong, K. and S. Zhang, Deciphering spatial domains from spatially resolved transcriptomics with an adaptive graph attention auto-encoder. Nat Commun, 2022. 13(1): p. 1739.

27. Hu, J., et al., SpaGCN: Integrating gene expression, spatial location and histology to identify spatial domains and spatially variable genes by graph convolutional network. Nat Methods, 2021. 18(11): p. 1342–1351.

28. Pham, D., et al., Robust mapping of spatiotemporal trajectories and cell-cell interactions in healthy and diseased tissues. Nat Commun, 2023. 14(1): p. 7739.

29. Chang, Y., et al., Define and visualize pathological architectures of human tissues from spatially resolved transcriptomics using deep learning. Comput Struct Biotechnol J, 2022. 20: p. 4600–4617.

30. Hao, Y., et al., Integrated analysis of multimodal single-cell data. Cell, 2021. 184(13): p. 3573–3587.e29.

31. Dries, R., et al., Giotto: a toolbox for integrative analysis and visualization of spatial expression data. Genome Biol, 2021.22(1): p. 78.

32. Zeira, R., et al., Alignment and integration of spatial transcriptomics data. Nat Methods, 2022. 19(5): p. 567–575.

33. Liu, X., R. Zeira, and B.J. Raphael, Partial alignment of multislice spatially resolved transcriptomics data. Genome Res, 2023. 33(7): p. 1124–1132.

34. Li, Z. and X. Zhou, BASS: multi-scale and multi-sample analysis enables accurate cell type clustering and spatial domain detection in spatial transcriptomic studies. Genome Biol, 2022.23(1): p. 168.

35. Guo, T., et al., SPIRAL: integrating and aligning spatially resolved transcriptomics data across different experiments, conditions, and technologies. Genome Biol, 2023.24(1): p. 241.

36. Liu, W., et al., Probabilistic embedding, clustering, and alignment for integrating spatial transcriptomics data with PRECAST. Nat Commun, 2023. 14(1): p. 296.

37. Stuart, T., et al., Comprehensive Integration of Single-Cell Data. Cell, 2019. 177(7): p. 1888–1902.e21.

38. Desai, O., et al., The Role of Immune and Inflammatory Cells in Idiopathic Pulmonary Fibrosis. Front Med (Lausanne), 2018. 5: p. 43.

39. Mages, S., et al., TACCO unifies annotation transfer and decomposition of cell identities for single-cell and spatial omics. Nat Biotechnol, 2023. 41(10): p. 1465–1473.

40. Chen, J., et al., ToppGene Suite for gene list enrichment analysis and candidate gene prioritization. Nucleic Acids Res, 2009. 37(Web Server issue): p. W305–11.

41. Avelar, R.A., et al., A multidimensional systems biology analysis of cellular senescence in aging and disease. Genome Biol, 2020.21(1): p. 91.

42. Saul, D., et al., A new gene set identifies senescent cells and predicts senescence-associated pathways across tissues. Nat Commun, 2022. 13(1): p. 4827.

43. Fridman, A.L. and M.A. Tainsky, Critical pathways in cellular senescence and immortalization revealed by gene expression profiling. Oncogene, 2008.27(46): p. 5975–87.

44. Casella, G., et al., Transcriptome signature of cellular senescence. Nucleic Acids Res, 2019. 47(14): p. 7294–7305.

45. De Cecco, M., et al., L1 drives IFN in senescent cells and promotes age-associated inflammation. Nature, 2019. 566(7742): p. 73–78.

46. Freund, A., C.K. Patil, and J. Campisi, p38MAPK is a novel DNA damage response-independent regulator of the senescence-associated secretory phenotype. Embo j, 2011. 30(8): p. 1536–48.

47. Besag, J., Spatial interaction and the statistical analysis of lattice systems. Journal of the Royal Statistical Society: Series B (Methodological), 1974. 36(2): p. 192–225.

48. Banerjee, S., B.P. Carlin, and A.E. Gelfand, Hierarchical modeling and analysis for spatial data. 2014: CRC press.

49. Gelman, A., et al., Bayesian data analysis. 2013: CRC press.

50. Eddelbuettel, D. and R. François, Rcpp: Seamless R and C++ integration. Journal of statistical software, 2011. 40: p. 1–18.

51. Chang, W., et al., shiny: Web Application Framework for R. R package version 1.7. 1. 2021.

