## Supplementary Materials for "Defining transcriptomic niches in human fibrotic lung with multi-sample spatial transcriptomics analysis using the MAPLE algorithm"

Hyeongseon Jeon *et al.*

**This PDF file includes:**

Figs. S1 to S3
Table S1

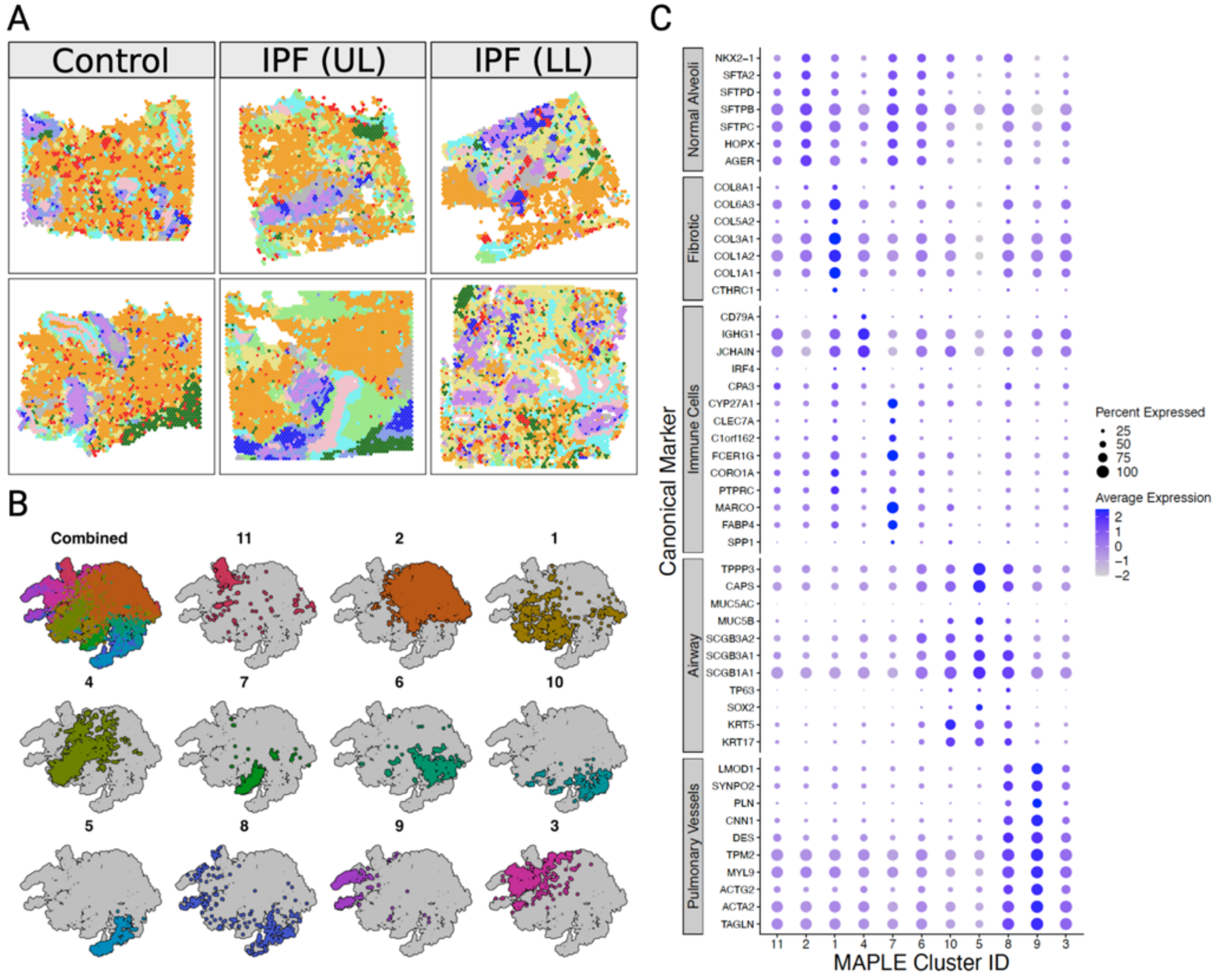


**Fig. S1.** MAPLE Clustering Prior to Pulmonary Region Annotation (**A**) Spatial distribution of 11 MAPLE clusters across six samples. (**B**) t-SNE visualization highlighting gene expression differences across MAPLE clusters. (**C**) Dot plot of canonical marker expression across MAPLE clusters.

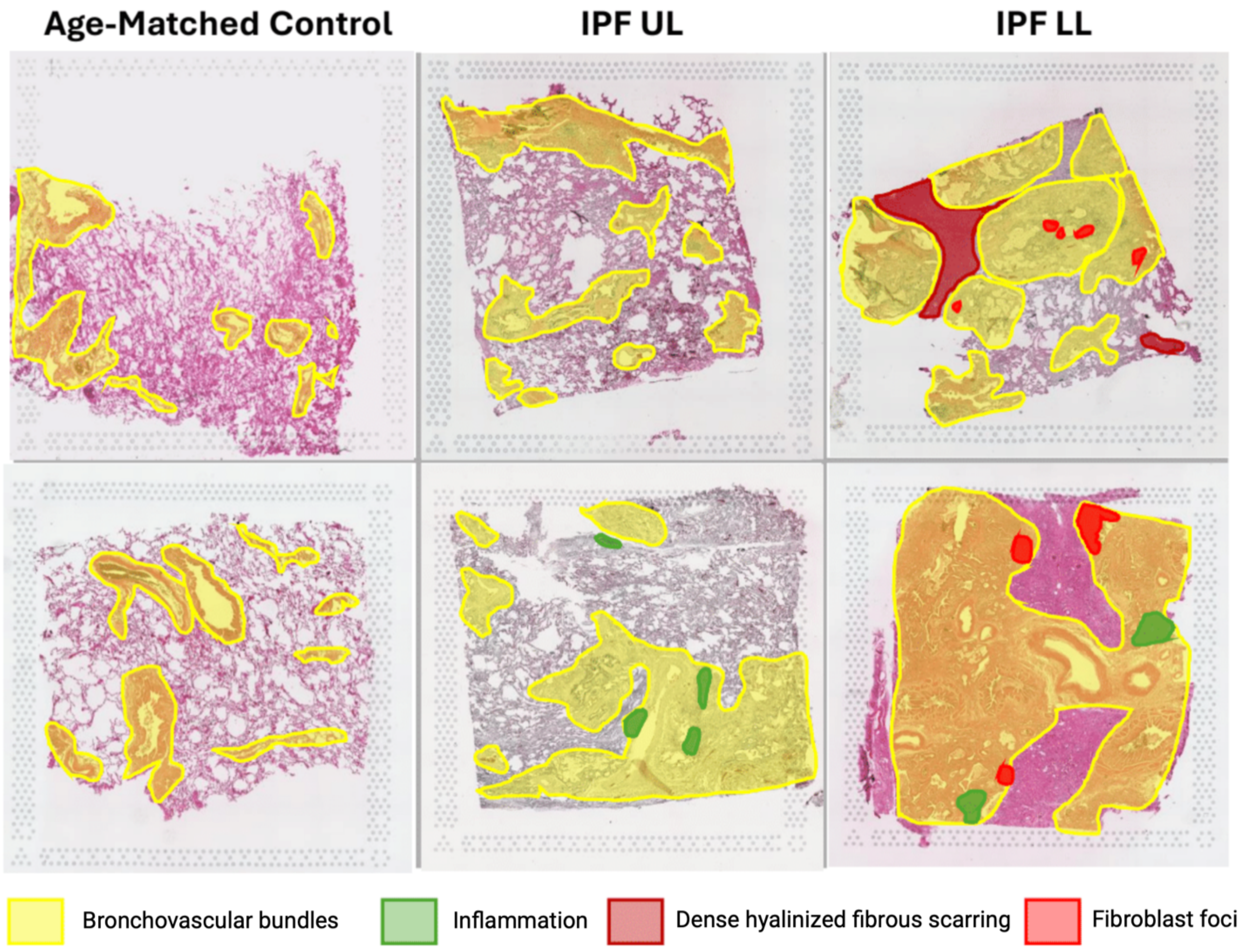


**Fig. S2. Overview of histopathologic annotations.** Yellow indicates bronchovascular bundles with surrounding interstitium, including lymphoid aggregates. Brown marks dense hyalinized fibrous scarring. Red denotes fibroblast foci. Green highlights inflammatory regions. Unmarked areas correspond to uninvolved or less-involved alveolar parenchyma.

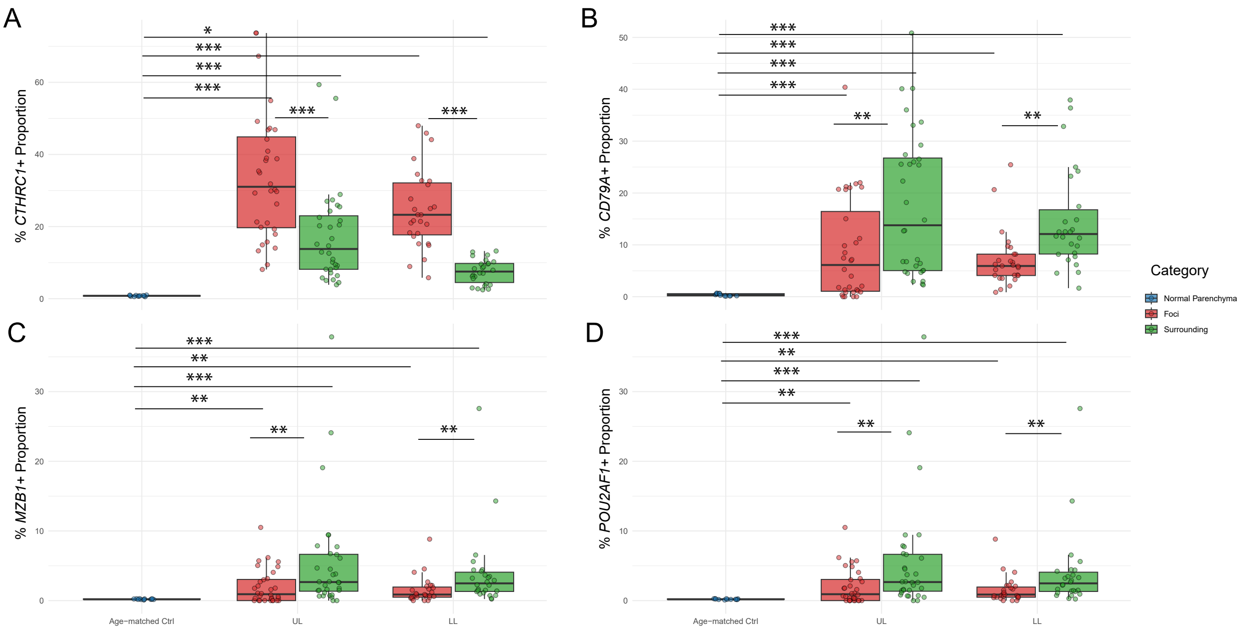


**Fig. S3.** 10 X Xenium spatial transcriptomics quantification of *CTHRC1* ⁺ fibroblasts and *CD79A* ⁺/*MZB1* ⁺/*POU2AF1* ⁺ B cells across fibrotic foci, surrounding regions, and age-matched parenchyma in IPF lung tissue sections.

**Table S1.** This table summarizes a stepwise gene-selection scheme. First, a conservatively curated list of SnC marker genes was assembled from multiple sources [41-46]. Second, Seurat was used to identify feature genes within this curated marker set. Finally, among these feature genes, only those showing positive expression patterns in lung-fibrosis samples were retained as the positively filtered subset.

| Description | Gene List |
|---|---|
| **Conservatively curated SnC marker gene list** | *CDKN1A,CDKN2A,EGFR,IGFBP1,IGFBP3,IL6,MIF,TP53,JUN,CDKN2B,IGFBP2,IGFBP5,IGFBP7,SERPINE1,ATM,AXL,ETS2,FAS,HRAS,IGF1,IGFBP6,IL15,IL7,LMNA,MAP2K3,MAP3K5,MAPK14,MAPK9,MDM2,MMP1,PARP1,PML,PRKCD,SIRT1,STAT1,TERF2,TERT,VEGFA,WNT16,WRN,ZMPSTE24,ANG,ARNTL,BCL2,BMI1,BRCA1,BUB1B,CALR,CBS,CCL1,CCL2,CCL26,CCNA2,CCND1,CDK1,CDK2,CDK6,CDKN1B,CEBPB,CLU,CTNNB1,CXCL1,CXCL8,CXCR2,E2F1,EEF1E1,ERBB2,EREG,FGF2,FGF7,FOS,FOXM1,FOXO1,GDF15,GSK3B,H2AFX,HDAC1,HELLS,HGF,HIF1A,HMGA1,HMGA2,HMGB1,HSPA1A,HSPA9,ICAM1,IFI16,IFNG,IGF1R,IGFBP4,IL13,IL1A,IL6ST,ING1,IRF5,IRF7,JAK2,KL,LMNB1,MAGEA2,MAP2K1,MAP2K6,MAP2K7,MAPK8,MAPKAPK5,MMP10,MMP12,MMP13,MMP14,MMP2,MMP3,MMP7,MMP9,MTOR,MYC,NBN,NEK6,NRG1,OPA1,PDGFB,PIGF,PIK3CA,PIN1,PLA2R1,PLAUR,PNPT1,PPARG,PPM1D,PRKDC,PTEN,RB1,RBL1,RBL2,RECQL4,RELA,SERPINB2,SIRT3,SIRT6,SMURF2,SOD1,SOD2,SP1,SRF,STAT3,TBX2,TBX3,TFDP1,TGFB1,TIMP2,TNF,TNFRSF10C,TNFRSF11B,TNFRSF1A,TOP1,TP63,TWIST1,TXN,ULK3,WNT2,YBX1,YPEL3* |
| **Seurat-derived feature genes within the curated SnC marker set** | *AXL,BUB1B,CCL2,CDKN1A,CDKN2B,CLU,CXCL8,ERBB2,EREG,FGF7,FOS,GDF15,ICAM1,IFNG,IGF1,IGFBP2,IGFBP3,IGFBP4,IGFBP5,IGFBP6,IGFBP7,IL6,JUN,KL,MMP10,MMP13,MMP14,MMP2,MMP7,MMP9,MYC,PDGFB,PLAUR,PPARG,SERPINE1,SOD2,TBX2,TBX3,TIMP2,TNF,TNFRSF11B,VEGFA,WNT2* |
| **Positively filtered genes from the feature-gene set in lung fibrosis** | *FGF7,GDF15,IGF1,IGFBP4,MMP13,MMP14,MMP2,MMP9,SERPINE1,TNF,WNT2* |
